# Motor learning is regulated by GDNF levels in postnatal cerebellar Purkinje cells

**DOI:** 10.1101/2024.09.06.611586

**Authors:** Elina Nagaeva, Giorgio Turconi, Kärt Mätlik, Mikael Segerstråle, Soophie Olfat, Vilma Iivanainen, Tomi Taira, Jaan-Olle Andressoo

## Abstract

Purkinje cells, the sole output neurons of the cerebellar cortex, are crucial for cerebellum-dependent motor learning. Previously we demonstrated that a ubiquitous 2-3-fold increase of endogenous glial cell line-derived neurotrophic factor (GDNF) improves motor learning. However, GDNF impacts many organ systems and cell types throughout the body leaving the underlying mechanism elusive. Here, we utilize an innovative conditional GDNF Hypermorphic mouse model to show that a 2-fold increase in endogenous GDNF specifically in postnatal Purkinje cells (PCs) is sufficient to enhance motor learning in adult animals. We demonstrate that improved motor learning is associated with increased glutamatergic input to PCs and elevated spontaneous firing rate of these cells, opposite to cerebellar ataxia where reduction in motor function and learning associates with decreased spontaneous activity of PCs. Notably, the GDNF expression levels variation range studied in our mouse model’s cerebellum falls within the normal range of variation observed in healthy human cerebellums. Our findings uncover a molecular pathway and a specific cell type that regulate motor learning, potentially explaining some individual differences in human motor skill acquisition.

## 1 INTRODUCTION

The cerebellum is a central regulator of motor function, including motor learning (De Zeeuw & Ten Brinke, 2015) and coordination (Manto et al, 2012). Purkinje cells (PCs) are the sole output source of signals that are integrated in the cerebellar cortex and are critical for dictating the cerebellum-dependent mechanisms regulating movements (Hirano, 2018; Witter et al, 2016). The stable firing rate of these relatively large inhibitory cells is necessary for efficient motor performance (Hosy et al, 2011), with downregulation linked to progressive deficits in motor coordination and learning in various forms of cerebellar ataxia (Hansen et al, 2013; Cook et al, 2021). Conversely, motor learning can be stimulated by direct optogenetic increase of PC firing rate (Nguyen-Vu et al, 2013; Lee et al, 2015; Silva et al, 2024). The fine-tuned activity of PCs relies on balanced interactions within the network of cerebellar cortex (Jelitai et al, 2016), where reduced excitatory inputs from granule cells or parallel fibres correlate with motor deficits in several mouse models (Bartelt et al, 2023; van der Heijden et al, 2023; Benedetti et al, 2016).

Glial cell line-derived neurotrophic factor (GDNF) is a secreted protein mostly known for exerting its trophic effect on dopamine neurons (Ibáñez & Andressoo, 2017; Hoffer et al, 1994; Lin et al, 1993). For this reason, ectopic delivery of GDNF has been tested in several preclinical and clinical studies of Parkinson’s disease, where striatal dopamine is lost due to the degeneration of nigrostriatal dopamine neurons (Whone et al, 2019a; Winkler et al, 1996; Gash et al, 1996; Sullivan et al, 1998; Beatty-Desana et al, 1975; Kordower et al, 2000; Whone et al, 2019b; Barker et al, 2020). We previously generated a mouse model where the replacement of the 3’ untranslated region (3’ UTR) of the *Gdnf* gene with a shorter sequence that does not contain binding sites for inhibitory molecules, such as microRNAs, increases the stability of Gdnf mRNA, resulting in a constitutive and ubiquitous about 2-fold elevation of GDNF production restricted to the cells that naturally transcribe Gdnf (GDNF Hypermorph mice, *Gdnf*^wt/Hyper^, Kumar et al, 2015). We found that *Gdnf*^wt/Hyper^ mice have improved motor function and motor learning, which lasts until old age (Mätlik et al, 2018; Turconi et al, 2020). Considering that GDNF is naturally expressed at several levels along the motor system, including its central (cortex, striatum, cerebellum) and peripheral parts (spinal cord, skeletal muscle tissue) (Henderson et al, 1994; Linnarsson et al, 2001; Golden et al, 1999; Sergaki et al, 2017; Hantman & Jessell, 2010; Hidalgo-Figueroa et al, 2012; Golden et al, 1998; Trupp et al, 1997), we wondered how the moderate 2-fold increase in GDNF levels elicit this phenotype and which organ system(s) or cell type(s) are involved.

In this study, we used a recently generated GDNF conditional Hypermorph (*Gdnf*^cHyper^) mouse model (Mätlik et al, 2022) in combination with Cre recombinase-based approaches and various knock-in reporter alleles to explore when, where, and how endogenous GDNF levels regulate motor learning. We also evaluated normal interindividual variation of GDNF expression levels in the human cerebellum to contextualize our findings.

## 2 MATERIALS AND METHODS

### 2.1 Animals

All animal experiments were carried out in accordance with the European Union Directive 86/609/EEC and were approved by the County Administrative Board of Southern Finland (license numbers ESAVI/11198/04.10.07/2014 and ESAVI/12046/04.10.07/2017). Mice were maintained in a 129Ola/ICR/C57bl6 mixed genetic background and in temperature-controlled conditions at 20°C– 22°C under a 12-h/12-h light/dark cycle at a relative humidity of 50%–60%. Cages and bedding material (Aspen chips; Tapvei Oy, Finland) were changed weekly, and wooden tube and aspen shavings were provided as enrichment. Mice received food and water *ad libitum*. The following mouse lines were used: *Gdnf*^Hyper^ (Kumar et al, 2015); *Gdnf*^cHyper^ (Mätlik et al, 2022); *Gdnf*^cKO^ (Kopra et al, 2015); *Pcp2*^Cre^ (Barski et al, 2000); *Ret*^GFP^ (Jain *et al*, 2006). All behavioural tests, tissue collection, processing, and analyses were performed by researchers blinded to the genotypes of the animals. For behavioural tests, only male mice were used (see below for more details), whereas other assessments were performed in both genders. In all experiments, wild-type littermates were used as controls.

### 2.2 Behavioural tests

All behavioural tests, including rotarod, vertical grid, open field, multiple static rods, coat hanger, beam walking, and grip strength, were performed as described in our previous studies (Mätlik *et al*, 2018; Turconi *et al*, 2020), following the protocols provided by the Laboratory Animal Centre of the University of Helsinki. Mice were tested at 2-3 months of age.

### 2.3 Stereotaxic surgeries

For intracranial surgeries, 2-month-old *Gdnf*^wt/wt^ and *Gdnf*^wt/cHyper^ mice were induced and maintained under anaesthesia with isoflurane (3-4% for induction and 1.5-2% for maintenance; Oriola, Finland) in 100% oxygen. After complete anaesthesia was achieved, the mice were placed on a stereotaxic apparatus and injected into the skin just above the skull with lidocaine for local analgesia (Yliopiston Apteekki, Finland). A 5µl Hamilton syringe with a 33-gauge needle (World Precision Instruments, USA) was used for intracranial injections. Mice were bilaterally injected into the striata with a recombinant AAV vector serotype 5 expressing Cre recombinase under the synapsin promoter (1.7 × 10^11^ viral genome copies) at the following coordinates relative to bregma at a 10-degree angle: A/P 1.2, M/L ±2.2; D/V: −3.2 mm. Viral vectors were produced in Lund University, Sweden, as previously described (Ulusoy *et al*, 2012). 1.5 μl of AAV5-Cre were injected into each site of the striatum at a flow rate of 0.2 μl/min. The needle was kept in place for an additional 5 minutes and retracted slowly. The skin was closed with sutures and animals were subcutaneously injected with 5 mg/kg of Rimadyl or Norocarp (Yliopiston Apteekki, Finland) in saline for postoperative analgesia.

### 2.4 Tissue isolation

For quantitative PCR, mice were euthanized by cervical dislocation followed by decapitation after deep anaesthesia with CO_2_. The brain was quickly removed from the skull, immersed in ice-cold 1X PBS, and the tissues of interest were dissected using a scalpel (*i.e.* cerebellum) or a puncher of 2 mm inner diameter (*i.e.* striatum), snap frozen, and stored at −80°C until processed.

For immunostaining, mice were anesthetized with pentobarbital (Mebunat, 200 mg/kg, i.p., Yliopiston Apteekki) and perfused with 1X PBS and 4% paraformaldehyde (PFA, Sigma). Brains were post-fixed in 4% PFA overnight at room temperature, dehydrated in 30% sucrose (Thermo Fisher Scientific) in 1X PBS at +4°C, and stored at −80°C prior to sectioning. Cerebellar tissues were sagittally cryosectioned at a thickness of 35 μm and stored in cryopreserving buffer containing 30% ethylene glycol (Fisher) and 20% glycerol (Acros Organics) in PBS at −20°C until processed.

### 2.5 RNA isolation and quantitative PCR

Total RNA was isolated from frozen tissues using Trizol Reagent (Thermo Fisher Scientific). For RNA analysis, 500 to 1000 ng of DNase I-treated total RNA was reverse transcribed to complementary DNA using random hexamer primers and RevertAid Reverse Transcriptase (Thermo Fisher Scientific). Complementary DNA was diluted 1:10 and stored at −20 °C until analysis. Quantitative PCR was performed with BioRad C1000 Touch Thermal Cycler upgraded to CFX384 System (BioRad), supplied with SYBR Green I Master (Roche) and 250 pmol primers, in 10 μl total volume in 384-well plates. Each sample was run in triplicate. The following primer pairs were used: *Gdnf* (F: 5’ CGCTGACCAGTGACTCCAATATGC 3’, R: 5’ TGCCGCTTGTTTATCTGGTGACC 3’), *Actb* (F: 5’ CTGTCGAGTCGCGTCCA 3’, R: 5’ ACGATGGAGGGGAATACAGC 3’), *Gapdh* (F: 5’ CCTCGTCCCGTAGACAAAA 3’, R: 5’ ATGAAGGGGTCGTTGATGGC 3’). The expression level of *Gdnf* was normalized to the geometric mean of *Actb* and *Gapdh* housekeeping gene expression. Results for a biological repeat were discarded when the Cq value for one or more of the replicates was 0 or above 40, or when the Cq difference between replicates was >1.

### 2.6 GDNF protein quantification

GDNF protein levels were measured with the GDNF Emax Immunoassay System (G7620, Promega), according to the protocol provided by the manufacturer. Mouse striatum samples were homogenized in a lysis buffer recommended by the manufacturer and the homogenates were centrifuged at 5000 rpm for 15 minutes at 4 °C. The supernatant was acid-treated using HCl, and GDNF protein levels and total protein levels were measured and analysed at the same to avoid freezing-thawing of the samples at any stage, as also described in (Kumar *et al*, 2015; Mätlik *et al*, 2022; Kopra *et al*, 2015). Similarly processed striatal samples from *Gdnf*^cKO^;Nestin-Cre mice(Kopra *et al*, 2015) were used as a negative control to set the background for GDNF protein measurements in all experiments.

### 2.7 Immunohistochemistry

Free-floating cerebellar sections were mounted on a SuperFrost Ultra Plus slides (Thermo Scientific) and air-dried overnight. Samples were washed 3×10 minutes in TBS and blocked for 1h in antibody solution (1X TBS containing 5% normal donkey serum and 0.1% Triton X-100). Primary antibodies were diluted in blocking solution (1X TBS containing 0.5% normal donkey serum and 0.1% Triton X-100) and samples were incubated with primary antibodies at +4°C overnight. After washing 3×10 minutes with 1X TBS, samples were incubated in darkness with secondary fluorescent antibodies in for 1h at room temperature, followed by 3×10 minutes washing with 1X TBS. Finally, the samples were mounted with ProLong Glass Antifade Mountant (Thermo Fisher) and kept at +4°C in the dark until imaging. The list of primary and secondary antibodies used in this study can be found in the Supplementary Table 1.

### 2.8 Image analysis

All fluorescence images were captured using ZEISS Axio Imager 2 wide-field microscope (Zeiss) and analysed with Fiji ImageJ version 1.53. All the analyses were conducted in the cerebellar lobe VIII in 2 months old *Ret*^GFP^;Pcp2-Cre;*Gdnf*^cHyper^ mice on two midsagittal images/animal, approximately at the same location, unless otherwise specified. Counts of RET-positive cells were performed in the cerebellar lobe VIII from four to six serial sections per animal (one section every 210 µm) containing all the layers of the cerebellar cortex. Similarly, the thickness of the cerebellar layers was analysed from the same images in two different locations of cerebellar lobe VIII, named A and B, as indicated in Figure S1C. The analysis of cerebellar sections stained with VGLUT2 and VGLUT1 antibodies was quantified using SynQuant, a Fiji ImageJ plugin(Wang *et al*, 2020), in a 200 µm^2^ region of interest (ROI) in the molecular layer of lobe VIII right above the Purkinje layer. More specifically, the total number of VGLUT2-positive puncta and total intensity of VGLUT1 staining were analysed in the ROI. The latter was done due to VGLUT1+ innervation in the molecular layer with now clear boundaries. The total intensity was then divided by the background to obtain a final optical density parameter (O.D). All confocal images were captured with Zeiss LSM 880 confocal microscope at 40x magnification.

### 2.9 Whole-cell patch-clamp recordings

Brains from P14-P21 *Gdnf*^Hyper^ mice were dissected in ice-cold (in mM) 124 NaCl, 3 KCl, 1.25 NaH_2_PO_4_, 10 MgSO_4_, 26 NaHCO_3_, 15 D-glucose, 1 CaCl_2_, saturated with 5 % CO_2_ / 95 % O_2_. Adult P60-P100 *Gdnf*^cHyper^;Pcp2-Cre mice were anaesthetized with Mebunate + Lidocaine mixture and brains were dissected in high-sucrose cutting solution, containing (in mM) 240 sucrose, 2.5 KCl, 1.25 Na_2_HPO_4_, 2 MgSO_4_, 1 CaCl_2_, 26 NaHCO_3_, and 10 D-Glucose (Voerman *et al*, 2022) to improve neuronal survival. Cerebellar slices (350 μm) were sagittally cut using a vibratome (Vibratome Co., St. Louis, MO, USA) in the above solutions and stored at room temperature in (in mM) 124 NaCl, 3 KCl, 1.25 NaH_2_PO_4_, 4 MgSO_4_, 26 NaHCO_3_, 15 D-glucose, 1 CaCl_2_, saturated with 5 % CO_2_ / 95 % O_2_. The slices were used 1–4 h after cutting. For electrophysiological recordings, the slices were placed in a submerged chamber and superfused with artificial cerebrospinal fluid (ACSF, in mM): 124 NaCl, 3 KCl, 1.25 NaH_2_PO_4_, 1 MgSO_4_, 26 NaHCO_3_, 15 D-glucose, 2 CaCl_2_, saturated with 5 % CO_2_ / 95 % O_2_, at a rate of 2–3 ml/min. Whole-cell recordings were obtained from Purkinje cells in cerebellar lobe VIII using the Multiclamp 700B amplifier (Molecular Devices, Sunnyvale, CA, USA). Cells were confirmed as Purkinje cells visually and by biocytin immunostaining. For measuring spontaneous currents in *Gdnf*^Hyper^ mice, Purkinje cells (PCs) were voltage-clamped at −50 mV with 4–5 MΩ pipettes filled with (in mM) 135 CsMeSO_4_, 10 Hepes, 0.5 EGTA, 4 Mg-ATP, 0.3 Na-GTP, 2 NaCl and biocytin 2mg/ml (285 mOsm), pH 7.2. At −60 mV, GABAergic sIPSCs were seen as outward and glutamatergic sEPSCs as inward. For the same experiment in adult *Gdnf*^cHyper^;Pcp2-Cre mice, the internal solution contained (in mM) 130 CsMeSO_4_, 10 Hepes, 0.5 EGTA, 4 Mg-ATP, 0.3 Na-GTP, 4 NaCl, 3 QX-314, 8 Na_2_-Phosphocreatine (290 mOsm), pH 7.2. Due to the higher post-synaptic activity in adult PCs, sIPSCs were recorded at −40 mV as outward currents and sEPSCs at - 75 mV as inward currents to separate them. Recordings where access resistance was higher than 25 MΩ were discarded. The analysis of post-synaptic currents was done in using the Mini Analysis Program 6.0.3. (Synaptosoft). The amplitude detection threshold was set between 10 and 25 pA (2–5 times the baseline root mean square noise level), and the detected events were verified visually. To analyze the frequency of spontaneous postsynaptic events, a total of 200 events were counted, beginning 30 seconds after the start of the recorded file. The frequency was then determined by dividing this event count by the time during which these events occurred. Spontaneous action potential firing was measured in on cell mode with pipettes containing extracellular solution. The frequency of action potentials were analysed in Clampfit 10.7 (Molecular Devices) by selecting 5 random 1-s-long intervals, counting number of events in each of them and calculating the mean.

### 2.10 Statistical analysis

In all the data sets, the normality of the data was determined with the Shapiro-Wilk test. Comparisons between two groups were analysed with Student or Welch’s t-test or Mann-Whitney U-test. Comparisons between three groups were performed using one-way or two-way Analysis of Variance (ANOVA) followed by Tukey’s or Sidak’s *post hoc* test. Outliers were identified with ROUT test (1%) and excluded from analysis. Statistical analyses were performed with PRISM (GraphPad software). Assessments with p < 0.05 were considered significant.

Fold change of the human cerebellum expression data was calculated according to the formula:

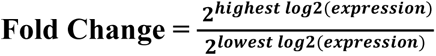

## 3 RESULTS

### 3.1 Increased levels of endogenous GDNF in the brain improve motor learning

Our previous analysis of constitutive *Gdnf*^Hyper^ mice (Kumar *et al*, 2015) revealed that *Gdnf*^wt/Hyper^ mice have improved motor performance, including motor learning in the rotarod and vertical grid tests (Mätlik *et al*, 2018; Turconi *et al*, 2020). However, motor function can be regulated by various central nervous system circuitries and peripheral tissues that express GDNF or its receptors. Next to the nigrostriatal dopamine system, these include the motor cortex, cerebellum, Clarke’s column of the spinal cord, dorsal root ganglia, and peripheral skeletal muscle tissue (Henderson *et al*, 1994; Linnarsson *et al*, 2001; Golden *et al*, 1999; Sergaki *et al*, 2017; Hantman & Jessell, 2010; Hidalgo-Figueroa *et al*, 2012; Golden *et al*, 1998; Trupp *et al*, 1997). Motor function and learning may be influenced by enhanced GDNF expression in all these sites, alone or in combination.

To study in which cell type and when GDNF improves motor function and learning, we used the GDNF <u>c</u>onditional <u>Hyper</u>morph mice (*Gdnf*^cHyper^) recently generated in our lab (Figure 1A, Mätlik et al, 2022). In these mice, the levels of endogenous GDNF can be increased at the post-transcriptional level in a spatiotemporally controlled manner via Cre-Lox-induced conditional 3’UTR replacement (Figure 1A, B). First, we crossed the *Gdnf*^cHyper^ mice with a Nestin-Cre line, where Cre recombinase activity is restricted to the CNS starting from about E12,5 (Tronche *et al*, 1999) (Figure 1B). Adult mice were tested with the rotarod test on two consecutive days, and the latency to fall off the rod was recorded over a period of 6 minutes. We found that *Gdnf*^wt/cHyper^;Nestin-Cre animals performed significantly better on the second day of the test compared to their littermate controls (Figure 1C), matching our previous observations in constitutive *Gdnf*^wt/Hyper^ mice (Mätlik *et al*, 2018; Turconi *et al*, 2020). Similarly, the time taken by *Gdnf*^wt/cHyper^;Nestin-Cre mice to turn from the down-facing to the up-facing position in the vertical grid test was significantly shorter on both days 1 and 2 compared with the control group (Figure 1D). These results indicate that improved motor performance and learning are induced by an increase in endogenous GDNF in the CNS after E12,5 when Cre starts to be expressed (Tronche *et al*, 1999), and allowed us to exclude an increase in GDNF in peripheral tissues as a significant contributor.

**Figure 1.**
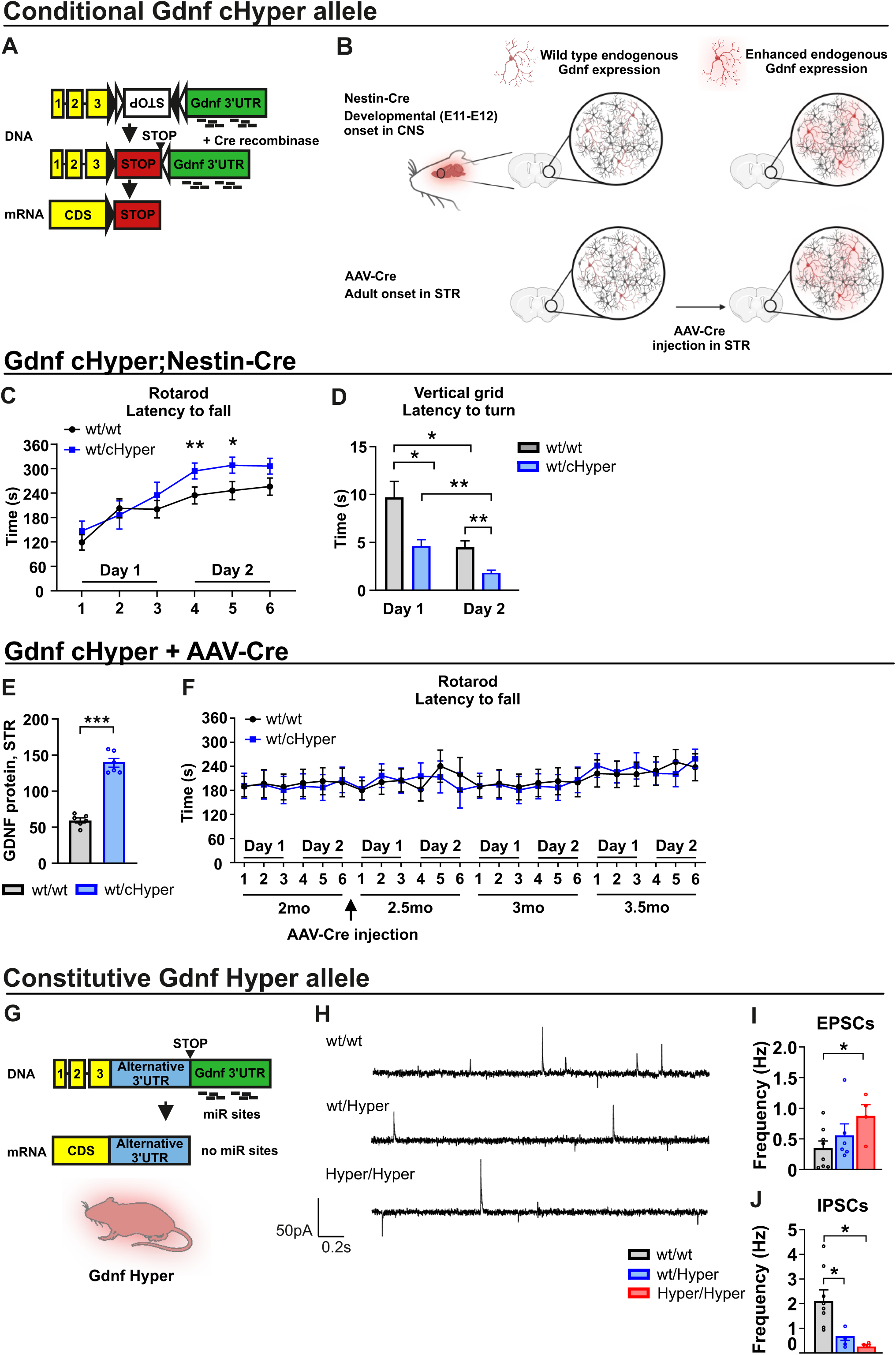
Brain, but not nigrostriatal, endogenous GDNF improves motor function and affects Purkinje cell postsynaptic currents. **(A)** Schematic of the *Gdnf*^cHyper^ allele. A cassette containing the bovine growth hormone polyadenylation signal (white box) was inserted in an inverted orientation flanked by the FLEx system immediately after the *Gdnf* stop codon. Upon Cre-mediated recombination, the FLEx cassette is inverted, and mRNA translation is terminated. The resulting Gdnf mRNA carries a shorter 3’UTR compared to native Gdnf 3’UTR (green box) which increases the stability of Gdnf mRNA due to the lack of microRNAs (miR) binding sites, thereby increasing GDNF production. **(B)** Elevated endogenous GDNF level in the CNS during development can be achieved upon crossing the *Gdnf*^cHyper^ allele with the Nestin-Cre mouse line (*Gdnf*^cHyper^;Nestin-Cre), whereas adult-onset increase of endogenous GDNF in the striatum is achieved by intrastriatal AAV-Cre injection in *Gdnf*^/cHyper^ mice (*Gdnf*^cHyper^ + AAV-Cre). **(C)** Latency to fall in the rotarod test and **(D)** latency to turn in the vertical grid test in *Gdnf*^wt/Hyper^;Nestine-Cre and their littermate controls at day 1 and day 2 of experiments. Unpaired t-test, Welch’s t-test, and Multiple t-test. *p < 0.05; **p < 0.05. **(E)** Striatal GDNF protein levels 2 months after bilateral striatal AAV-Cre injection of *Gdnf*^wt/wt^ and *Gdnf*^wt/cHyper^ mice. Welch’s t-test, ***p < 0.001; **(F)** Motor coordination and learning evaluated with the rotarod test on two consecutive days (Day 1 and Day 2), measured at baseline (before AAV-Cre injections) and at 2, 5, and 7 weeks after intrastriatal bilateral AAV-Cre injection in *Gdnf*^wt/wt^ and *Gdnf*^wt/cHyper^ mice. Multiple t-tests. **(G)** Schematic of the *Gdnf*^Hyper^ allele. The Pgk1-puΔtk-bGHpA sequence (alternative 3’UTR, blue box) was inserted immediately after the *Gdnf* stop codon. This cassette does not contain binding sites for inhibitory molecules, *i.e.* miR, compared to the Gdnf native 3’UTR (green box), resulting in increased Gdnf mRNA stability. **(H)** Sample traces of spontaneous excitatory and inhibitory postsynaptic currents (EPSCs and IPSCs, respectively) in *Gdnf*^Hyper^ mice and littermate controls. EPSCs are seen as inward currents (downward deflections) and IPSCs are seen as outward currents (upward deflections). **(I)** Spontaneous EPSCs and **(J)** IPSCs in Purkinje cells in cerebellar lobe VIII in *Gdnf*^Hyper^ mice. One-way ANOVA followed by Tukey’s multiple comparisons test. *p < 0.05. **(C, D)** *Gdnf*^wt/wt^;Nestin-Cre (n=12-23); *Gdnf*^wt/cHyper^;Nestin-Cre (n=8-10). **(E)** *Gdnf*^wt/wt^ (n=6); *Gdnf*^wt/cHyper^ (n=6). **(F)** *Gdnf*^wt/wt^ (n=15); *Gdnf*^wt/cHyper^ (n=14). **(I, J)** *Gdnf*^wt/wt^ (n=8); *Gdnf*^wt/Hyper^ (n=6); *Gdnf*^Hyper/Hyper^ (n=4). Data are presented as mean ± SEM. Figure created with BioRender.com.

In the nigrostriatal tract, GDNF is almost exclusively expressed in a highly specific neuron type, the parvalbumin-positive interneurons in the striatum (Hidalgo-Figueroa *et al*, 2012). GDNF is a strong trophic factor for dopamine neurons in the nigrostriatal pathway, the dopaminergic tract that supports motor function and degenerates in Parkinson’s disease (Hoffer *et al*, 1994; Quintino *et al*, 2019; Bondarenko & Saarma, 2021). Therefore, we first tested if increased endogenous GDNF in the nigrostriatal system improves motor function and learning. To that end, we injected AAV-Cre into the striata (Figure 1B) of young adult *Gdnf*^wt/cHyper^ mice and *Gdnf*^wt/wt^ controls to evaluate the contribution of increased endogenous GDNF in the nigrostriatal pathway on motor function with the rotarod test. Two months after AAV-Cre injection when adult-onset change in striatal endogenous GDNF levels is known to enhance dopamine function (Mätlik *et al*, 2022) and motor activity (Kopra *et al*, 2017), we found a 2-3-fold increase in striatal GDNF protein expression (Figure 1E), as expected based on earlier work (Mätlik et al., 2022). Despite increased GDNF expression, however, we did not observe differences in the rotarod performance, evaluated at baseline and at 2, 5, and 7 weeks after AAV-Cre injections (Figure 1F). These results suggest that an adult-onset increase in endogenous GDNF in the nigrostriatal pathway does not improve motor function and learning.

The observed superior performance in the vertical grid test and accelerating rotarod test may also relate to cerebellum and spinocerebellar tract function (Boisgontier and Swinnen, 2014), where GDNF is expressed in Purkinje cells (Sergaki et al., 2017), some granule neurons (Golden et al., 1999, 1998; Sergaki et al., 2017; Trupp et al., 1997), and neurons of the Clarke’s column in the spinal cord which project into cerebellar lobe VII and VIII (Hantman and Jessell, 2010). Since Purkinje cells integrate and provide the output of the cerebellar cortex (Hirano, 2018), we first studied the synaptic inputs converging onto Purkinje cells by using whole-cell patch-clamp recordings in cerebellar slices in constitutive *Gdnf*^Hyper^ mice (Figure 1G). We recorded spontaneous excitatory and inhibitory postsynaptic currents (EPSCs and IPSCs, respectively) in the somatic area of Purkinje cells in the cerebellar lobe VIII between postnatal days (P) 14-21. We found that the frequency of EPSCs was significantly higher in *Gdnf*^Hyper/Hyper^ mice compared to their control littermates (Figure 1H, I), whereas the frequency of IPSCs was lower in both *Gdnf*^wt/Hyper^ and *Gdnf*^Hyper/Hyper^ animals (Figure 1H, J). These findings demonstrated that increased GDNF levels in the constitutive *Gdnf*^Hyper^ mice result in changes in Purkinje cell synaptic responses. Unfortunately, it is not possible to analyse motor function and learning in adult *Gdnf*^Hyper/Hyper^ animals, since about 50% of mice die due to malformed urogenital block by P7 and 100% by P25 (Kumar et al., 2015). Therefore, we utilised a conditional approach to investigate the role of GDNF upregulation in Purkinje cells.

### 3.2 Postnatal Purkinje cell specific increase of endogenous GDNF level in *Gdnf*^cHyper^;Pcp2-Cre mice

The observed changes in EPSC/IPSC balance in Purkinje cells in both constitutive *Gdnf*^wt/Hyper^ and *Gdnf*^Hyper/Hyper^ animals prompted us to study the outcome of increased endogenous GDNF specifically in postnatal cerebellar Purkinje cells using the *Gdnf*^cHyper^ allele in combination with the Purkinje cell-protein 2 promoter driven Cre line (*Gdnf*^cHyper^;Pcp2-Cre) (Figure 2A). In *Gdnf*^cHyper^;Pcp2-Cre mice the activity of Cre recombinase is restricted to Purkinje cells in the cerebellum and retinal bipolar neurons starting from the first postnatal week (Sługocka et al., 2017).

**Figure 2.**
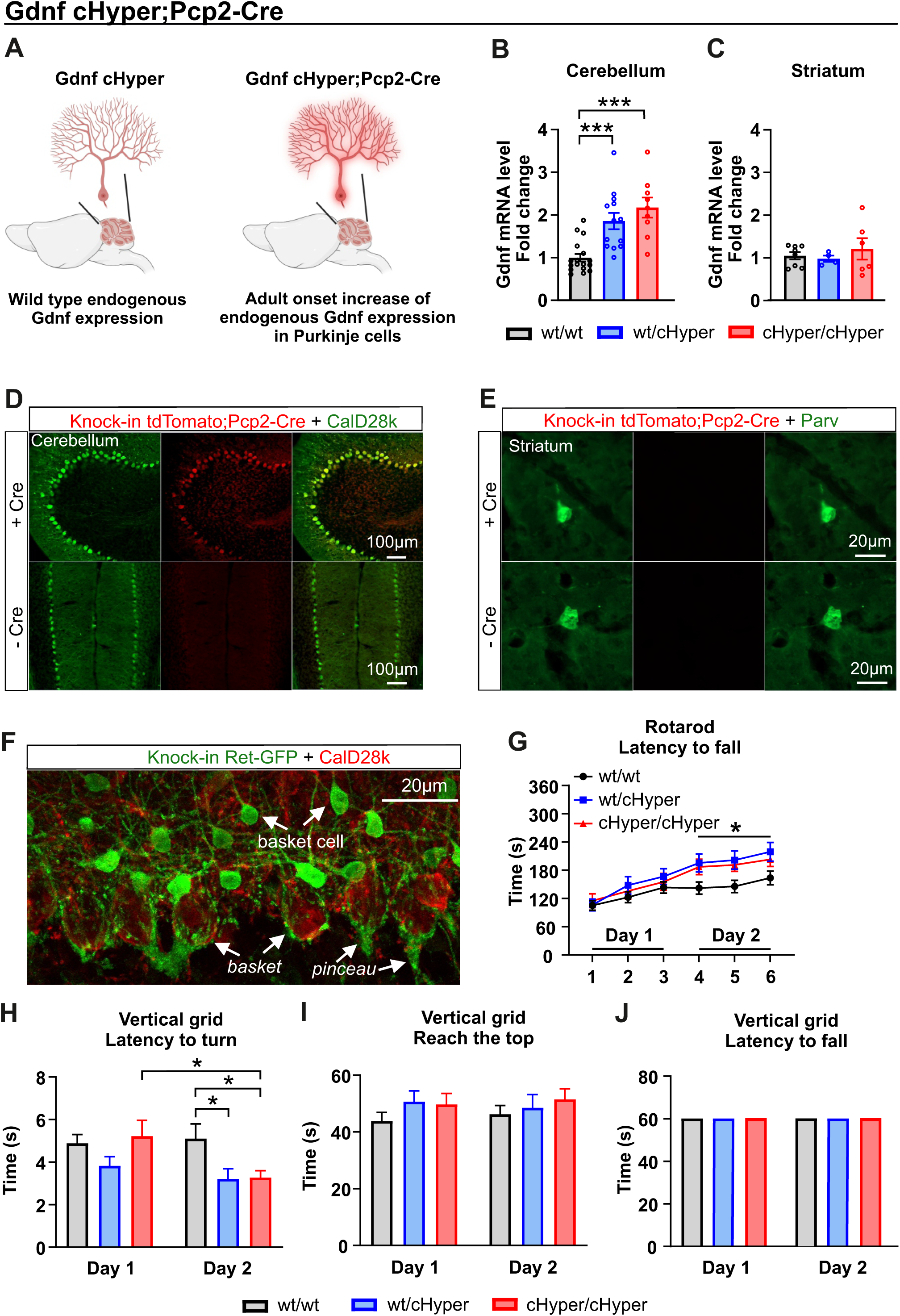
Increased endogenous GDNF level in Purkinje cells improves motor learning. **(A)** Increased endogenous GDNF in cerebellar Purkinje cells was achieved by crossing the *Gdnf*^cHyper^ allele with the Pcp2-Cre mouse line (*Gdnf*^cHyper^;Pcp2-Cre). **(B)** Cerebellar and (**C**) striatal Gdnf levels in adult *Gdnf*^cHyper^;Pcp2-Cre mice measured with qPCR. One-way ANOVA followed by Tukey’s multiple comparisons test. ***p < 0.001. **(D)** Representative cerebellar sagittal sections (upper and lower left panels) and (**E**) striatal coronal sections (upper and lower right panels) from adult tdTomato;Pcp2-Cre mice showing tdTomato signal (red) and calbindin (CalD28k, a marker for cerebellar Purkinje cells, left panel) or parvalbumin (Parv, marker for striatal GDNF-expressing parvalbumin interneurons, right panel). Scale bars are indicated in the figures. **(F)** Confocal image of a representative cerebellar sagittal section from an adult *Ret*^GFP^ mouse showing GFP signal in green and calbindin (CalD28k) in red. The receptor for GDNF, RET, is expressed in basket cell interneurons which extend their axon collaterals and form the *basket* around the Purkinje cell somata and the *pinceau* around the Purkinje cells axon initial segment. Scale bars are indicated in the figures. **(G)** Latency to fall in the rotarod test in *Gdnf*^cHyper^;Pcp2-Cre mice and wild-type littermates. Two-way ANOVA followed by Bonferroni’s multiple comparison test, genotype effect at Day 1 and Day 2. *p < 0.05. **(H)** Latency to turn, **(I)** latency to reach the top and (**J**) to fall of the grid in the vertical grid test in *Gdnf*^cHyper^;Pcp2-Cre mice and control group (wt/wt). Unpaired t-test and one-way ANOVA followed by Tukey’s multiple comparisons test. *p < 0.05. **(B)** *Gdnf*^wt/wt^;Pcp2-Cre (n=17); *Gdnf*^wt/cHyper^;Pcp2-Cre (n=13); *Gdnf*^cHyper/cHyper^;Pcp2-Cre (n=9). (**C**) *Gdnf*^wt/wt^;Pcp2-Cre (n=8); *Gdnf*^wt/cHyper^;Pcp2-Cre (n=4); *Gdnf*^cHyper/cHyper^;Pcp2-Cre (n=6). **(G-J)** *Gdnf*^wt/wt^;Pcp2-Cre (n=42-44); *Gdnf*^wt/cHyper^;Pcp2-Cre (n=22-23); *Gdnf*^cHyper/cHyper^;Pcp2-Cre (n=18-20). Data are presented as mean ± SEM. Figure created with BioRender.com.

First, we measured Gdnf expression in the cerebellum of *Gdnf*^cHyper^;Pcp2-Cre mice and found that Gdnf levels are significantly increased in both *Gdnf*^wt/cHyper^;Pcp2-Cre and *Gdnf*^cHyper/cHyper^;Pcp2-Cre mice compared with controls by about 1,9 and 2,2-fold respectively (Figure 2B). In the striatum, the brain region where GDNF is highly expressed (Hidalgo-Figueroa et al., 2012; Kumar et al., 2015), the expression level of Gdnf was comparable between genotypes (Figure 2C), demonstrating that, as expected, Pcp2-Cre does not affect GDNF expression levels in the striatum (Sługocka et al., 2017). Interestingly, despite the trend Gdnf expression in the cerebellum did not statistically significantly differ between heterozygotes and homozygotes, suggesting a possible ceiling effect for Gdnf expression in Purkinje cells (Figure 2B). To further assess the specificity of Pcp2-Cre recombination in the cerebellum, we crossed Pcp2-Cre mice to lox-STOP-lox tdTomato reporter mice. Immunohistochemical analysis of cerebellum from lox-STOP-lox-tdTomato;Pcp2-Cre mice revealed that Cre expression is indeed confined to cerebellar Purkinje cells and co-express with calbindin (CalD28k), a well-known Purkinje cell marker (Figure 2D, Barski et al, 2003). In line with our qPCR results in the striatum (Figure 2C), no Cre-expressing cells were found in striatal GDNF-producing parvalbumin GABAergic interneurons (Figure 2E, Hidalgo-Figueroa et al., 2012).

Next, we asked which cells in the cerebellum express the receptor tyrosine kinase RET, a signalling receptor for GDNF believed to mediate most cell-type-specific effects of GDNF (Airaksinen et al., 1999; Drinkut et al., 2016; Ibáñez and Andressoo, 2017; Taraviras et al., 1999). Earlier studies using in situ hybridization revealed Ret expression in the Purkinje cell layer of the cerebellar cortex (Golden et al., 1998; Trupp et al., 1997), but the exact cell type has remained unclear. To avoid artifacts related to immunohistochemical detection of RET, a commonly acknowledged limitation in the field, we utilized a *Ret*^GFP^ knock-in mouse line (Jain et al., 2006). Analysis of *Ret*^GFP^ knock-in mice revealed that RET is expressed in the basket cells, which are located in close proximity to the Purkinje cells in the Purkinje cell layer (Figure 2F), but not in Purkinje cells. Basket cells are inhibitory interneurons that support and regulate Purkinje cell function and development by forming the *basket* around Purkinje cell somata and the *pinceau* at the Purkinje cell axon initial segment (Figure 2F, Brown et al., 2019; Hirano, 2018).

To confirm that increased GDNF do not alter gross cerebellar anatomy, we first determined the number of RET-expressing basket cells. We used the *Ret*^GFP^ mouse line crossed with *Gdnf*^cHyper^;Pcp2-Cre mice to count the number of RET-expressing basket cells (Figure 2F), and found no difference between *Gdnf*^wt/wt^;Pcp2-Cre, *Gdnf*^wt/cHyper^;Pcp2-Cre, and *Gdnf*^cHyper/cHyper^;Pcp2-Cre mice (Figure S1A). Next, we measured the thickness of the three layers of the cerebellar cortex: the granule cell layer, the Purkinje cell layer, and the molecular layer, as indicated in Figure S1C. We observed no difference between the genotypes in any of the analysed layers (Figure S1D-F). These results suggested that a postnatal increase in endogenous GDNF in Purkinje cells does not alter the number of RET-expressing basket cells or the thickness of the cerebellar cortical layers.

### 3.3 Increase in endogenous GDNF in postnatal Purkinje cells increases motor learning

We next asked if increased levels of endogenous GDNF specifically in Purkinje cells regulate motor learning. To that end, we analysed *Gdnf*^cHyper^;Pcp2-Cre mice with the same behavioural tests previously carried out in constitutive *Gdnf*^wt/Hyper^ animals (Kumar et al., 2015; Mätlik et al., 2018; Turconi et al., 2020). We performed the accelerating rotarod test with three trials per day on two consecutive days. The latency to fall off the rod was comparable between groups in all three trials during the first day of the test (day 1) (Figure 2G). On day 2, however, *Gdnf*^wt/cHyper^;Pcp2-Cre and *Gdnf*^cHyper/cHyper^;Pcp2-Cre mice performed significantly better compared to the control group (Figure 2G), suggesting improved motor learning in mice with elevated GDNF expression in Purkinje cells. Next, we conducted the vertical grid test. The time to turn upward, to climb to the upper edge and to fall off the grid were measured with one trial per day on two consecutive days. The latency to turn on the grid was comparable between genotypes on the first day of the test (Figure 2H). Similarly to the rotarod test, however, *Gdnf*^wt/cHyper^;Pcp2-Cre and *Gdnf*^cHyper/cHyper^;Pcp2-Cre mice turned significantly faster compared to the control group on the second day (Figure 2H). We found no difference in the time to reach the top of the grid (Figure 2I) and none of the mice fell off the grid during the experiment (Figure 2J) reflecting no changes in the grip strength or basic motor performance. We conclude that a 2-fold increase in GDNF levels in postnatal Purkinje cells is sufficient to induce improved motor learning.

Next, we asked if and how the loss of Purkinje cell specific GDNF impacts motor function. For this, we crossed the Pcp2-Cre mice with mice carrying a conditional Knock-Out allele for Gdnf (*Gdnf*^cKO^) (Figure S2A, Kopra et al., 2015). We first analysed Gdnf levels in the cerebellum of *Gdnf*^cKO^;Nestin-Cre and *Gdnf*^cKO/KO^;Pcp2-Cre mice. As reported previously (Kopra et al., 2015), we observed full loss of Gdnf expression in the cerebellum (Figure S2B) of *Gdnf*^cKO^;Nestin-Cre mice. In *Gdnf*^cKO/KO^;Pcp2-Cre mice, we observed about 50% reduction in *Gdnf* mRNA levels in the cerebellum, indicating that the remaining cerebellar Gdnf mRNA is produced in other cell types than Purkinje cells, such as granule neurons, which have also been reported (Sergaki et al, 2017). Next, we analysed motor function in *Gdnf*^cKO^;Pcp2-Cre mice. We found no differences in the rotarod nor the vertical grid tests (Figure S2C, D). Altogether, these data suggest that an increase in endogenous GDNF in postnatal Purkinje cells is sufficient to improve motor learning, whereas the deletion of GDNF in these neurons has no effect on motor function.

### 3.4 Increased endogenous GDNF expression in Purkinje cells does not affect voluntary motor behaviour and muscle strength

Next, we performed additional tests to further assess motor behaviour. We first evaluated spontaneous locomotor activity for 30 minutes with the open field test and found no differences in the total distance travelled nor the time spent in the centre of the open field arena between groups (Figure 3A, B). Analysis with the multiple static rods test, which consist of five rods of decreasing diameter revealed that the latency to turn on each rod and the time to reach the platform were comparable between genotypes (Figure 3C, D). Next, motor balance was evaluated with the beam walking test. The number of sections crossed and the latency to fall off the beam were not different between genotypes (Figure 3E, F). Because muscle strength may affect motor function, we analysed this parameter with the coat hanger and grip strength tests but found no difference between genotypes (Figures 3H, G; see Materials and Methods for details). Collectively, these results indicate that GDNF levels in postnatal Purkinje cells regulate motor learning in tests which require immediate reaction and motor response, but do not impact motor performance in tests where animals can progress slowly on the voluntary basis. In addition, these results show that GDNF levels in Purkinje cells do not impact muscle strength.

**Figure 3.**
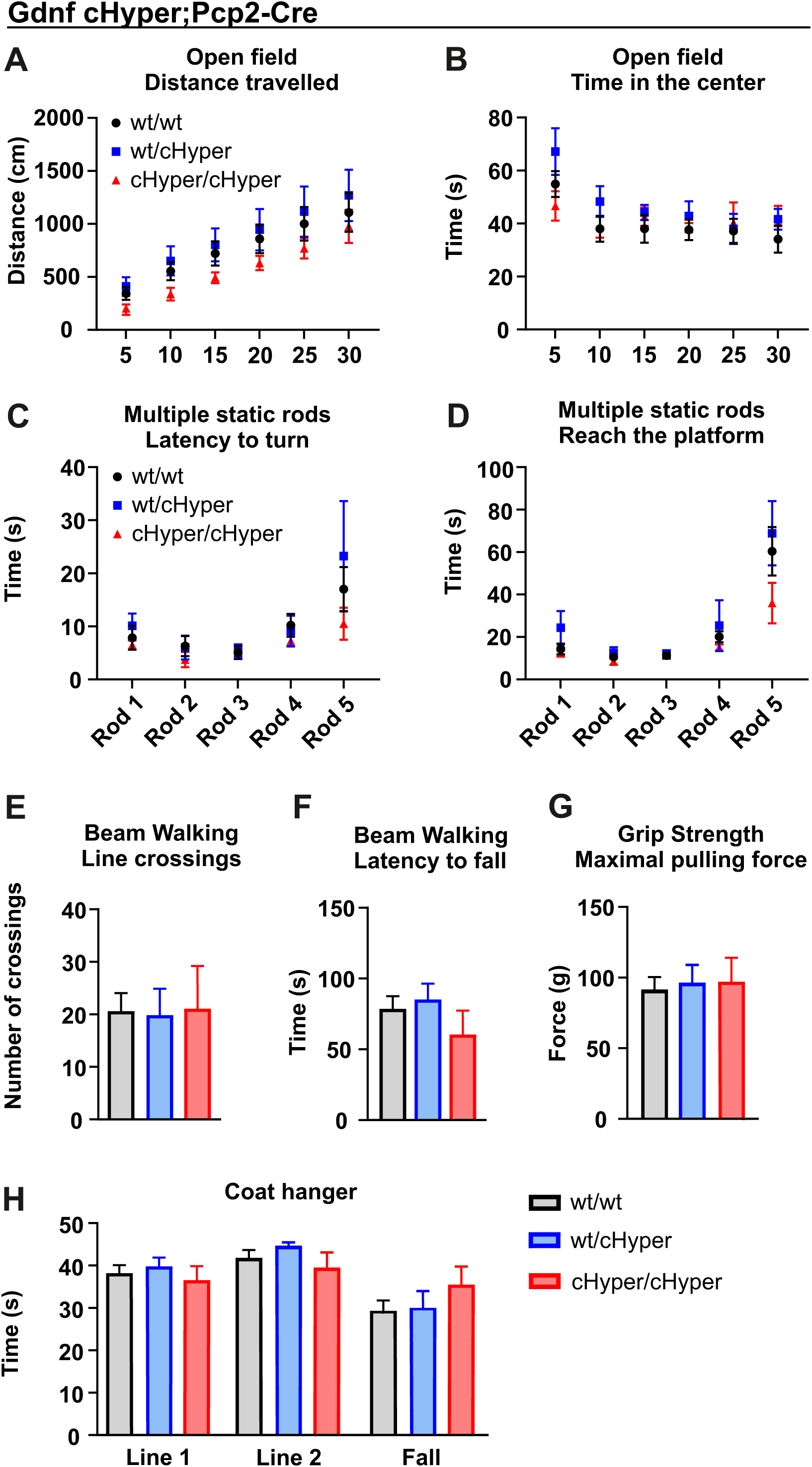
Unaltered voluntary motor behaviour and muscle strength in *Gdnf*^cHyper^;Pcp2-Cre mice. **(A)** Total distance travelled and **(B)** time spent in the centre of the arena in the open field test calculated in 5-minute blocks. Two-way ANOVA followed by Bonferroni’s multiple comparison test. **(C)** Latency to turn and **(D)** time to reach the platform on each rod in the multiple static rods test. Two-way ANOVA followed by Bonferroni’s multiple comparison test. **(E)** Number of crossed lines and **(F)** latency to fall from the beam during beam walking test. One-way ANOVA followed by Tukey’s multiple comparisons test. **(G)** Maximal forepaw pulling force measured in the grip strength test. One-way ANOVA followed by Tukey’s multiple comparisons test. **(H)** Latency to reach the lines (Line 1: end of horizontal part; Line 2; diagonal part of the coat hanger) and latency to fall in the coat hanger test. One-way ANOVA followed by Tukey’s multiple comparisons test. Data are presented as mean ± SEM. **(A-D)** *Gdnf*^wt/wt^;Pcp2-Cre (n=14-38); *Gdnf*^wt/cHyper^;Pcp2-Cre (n=9-20); *Gdnf*^cHyper/cHyper^;Pcp2-Cre (n=4-13).

### 3.5 Increased endogenous GDNF levels alter postsynaptic inputs and firing frequency of Purkinje Cells

Given that we observed increased EPSC frequency and decreased IPSC frequency in Purkinje cells upon a ubiquitous and constitutive increase of endogenous GDNF levels in P14-P21 *Gdnf*^Hyper^ mice (Figure 1H-J), we measured the same parameters in adult *Gdnf*^cHyper^;Pcp2-Cre mice to temporally correspond to behavioural experiments which were performed in young adult animals (at 2-4-months of age, termed P60-P100). We performed whole-cell patch-clamp recordings in Purkinje cell somata in young adult *Gdnf*^cHyper/cHyper^;Pcp2-Cre mice and littermate controls, and found that the frequency of spontaneous EPSCs was increased (Figure 4A, B), whereas the frequency of spontaneous IPSCs in *Gdnf*^cHyper/cHyper^;Pcp2-Cre mice was not changed compared to their littermate controls (Figure S3A-B). To better understand the mechanisms underlying altered postsynaptic currents in Purkinje cells, we quantified the excitatory synapses onto Purkinje cells in *Gdnf*^cHyper/cHyper^;Pcp2-Cre mice using immunohistochemistry. Purkinje cells receive two major excitatory synaptic inputs: climbing fibres, originating from the inferior olivary complex, which use the vesicular glutamate transporter 2 (VGLUT2, Hioki et al., 2003), and parallel fibres, originating from the cerebellar granule cells, which predominantly use VGLUT1 in adult animals (Miyazaki et al., 2003). We found an increase in the VGLUT2-positive glutamatergic inputs onto Purkinje cells in *Gdnf*^cHyper/cHyper^;Pcp2-Cre mice relative to the wild type controls (Figure 4D). Similarly, the VGLUT1 immunoreactivity was enhanced in *Gdnf*^cHyper/cHyper^;Pcp2-Cre mice as was calculated by comparing the optical density between genotypes (Figure 4C). These results suggest that GDNF elevation in *Gdnf*^cHyper/cHyper^;Pcp2-Cre mice increases the number of excitatory synapses on Purkinje cells.

**Figure 4.**
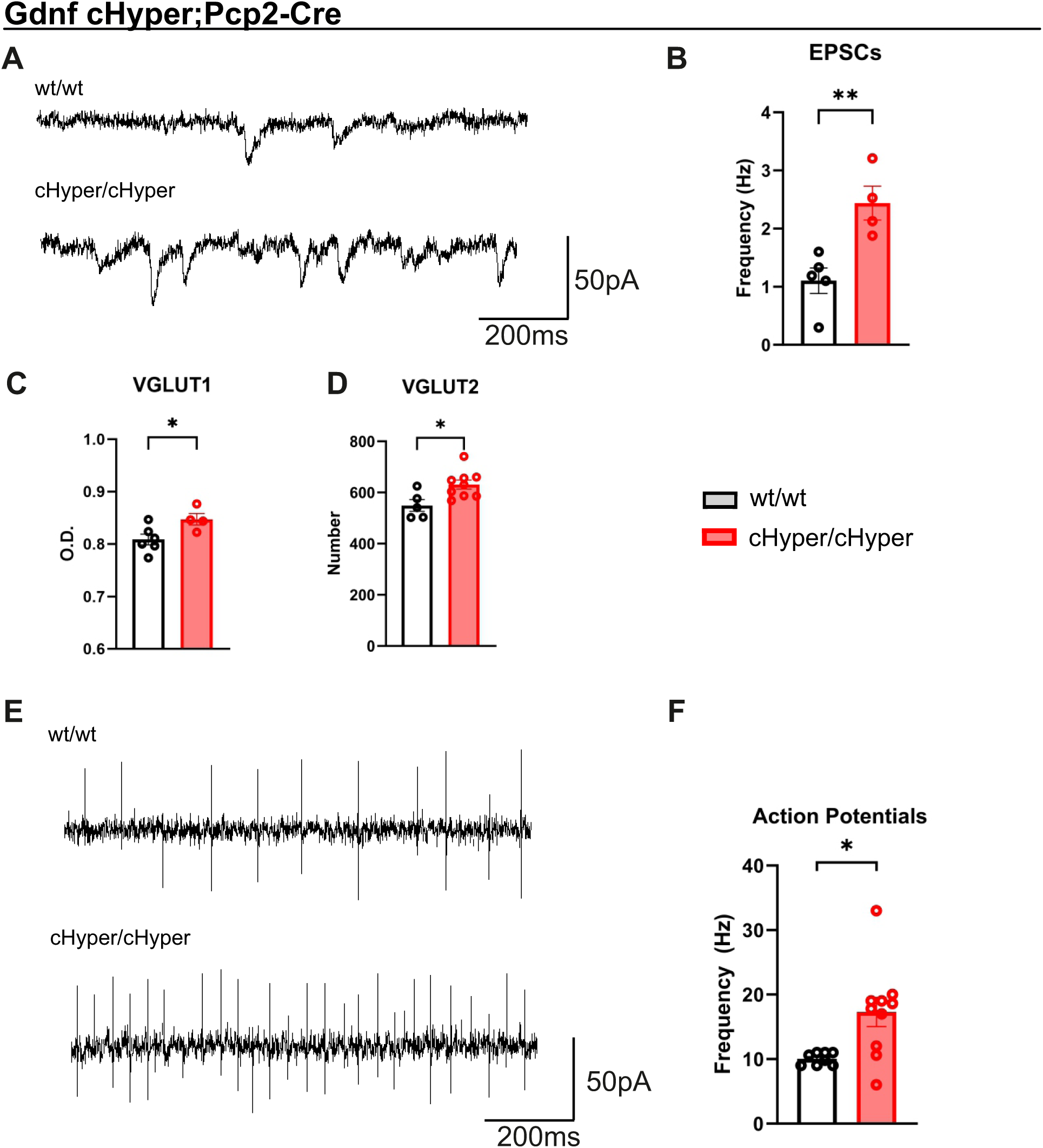
Changes in Purkinje cell synaptic responses and firing frequency in *Gdnf*^cHyper^;Pcp2-Cre mice. **(A)** Sample traces of spontaneous excitatory post synaptic currents (EPSCs) in *Gdnf*^cHyper/cHyper^;Pcp2-Cre mice and their littermate controls. EPSCs are seen as inward currents (downward deflections) at the −75 mV holding potential. **(B)** Spontaneous EPSC frequency in Purkinje cells measured in cerebellar lobe VIII of *Gdnf*^wt/wt^;Pcp2-Cre adult mice and littermate controls. Unpaired t-test. ** p < 0.01. **(C)** Optical density (O.D.) of vesicular glutamate transporter 1 (VGLUT1) and **(D)** number of vesicular glutamate transporter 2 (VGLUT2) synapses in 200 µm^2^ area within the molecular layer in the proximity to PC layer. Unpaired t-test. *p < 0.05. **(E)** Sample traces from on cell recordings and **(F)** action potential firing frequency from Purkinje cells in controls (wt/wt) and *Gdnf*^cHyper/cHyper^;Pcp2-Cre mice. Unpaired t-test. *p < 0.05. Data are presented as mean ± SEM. **(B)** *Gdnf*^wt/wt^;Pcp2-Cre (n=5); *Gdnf*^cHyper/cHyper^;Pcp2-Cre (n=4). **(C)** *Gdnf*^wt/wt^;Pcp2-Cre (n=6); *Gdnf*^wt/cHyper^;Pcp2-Cre (n=4); **(D)** *Gdnf*^wt/wt^;Pcp2-Cre (n=5); *Gdnf*^wt/cHyper^;Pcp2-Cre (n=8);. **(F)** *Gdnf*^wt/wt^;Pcp2-Cre (n=8); *Gdnf*^cHyper/cHyper^;Pcp2-Cre (n=10).

**Figure 5.**
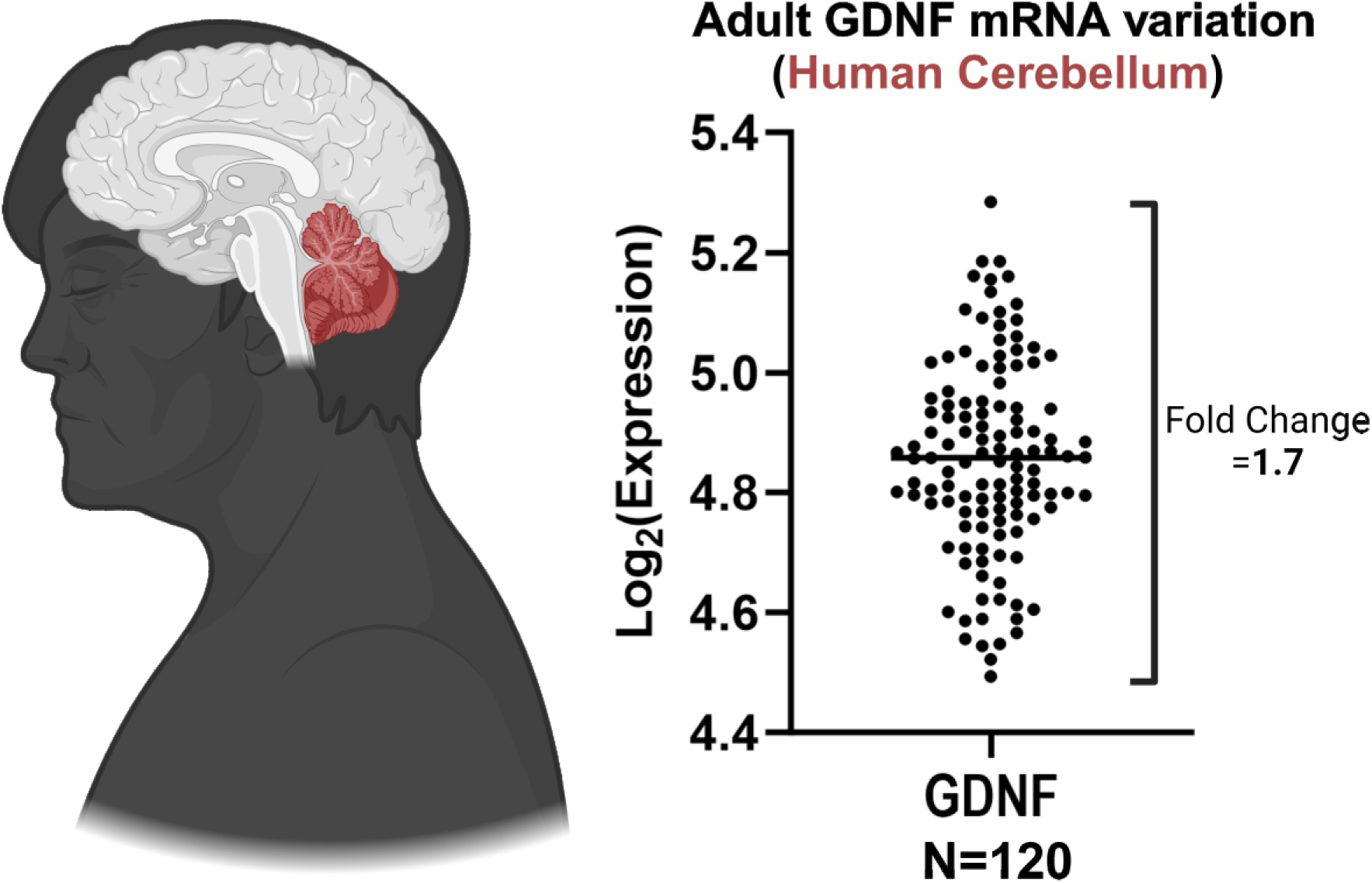
Distribution of GDNF expression levels in the healthy human cerebellum. Expression levels of GDNF mRNA in the cerebellum of healthy human donors, detected using the whole transcriptome Affymetrix Assay and published earlier (Ramasamy et al., 2014). The Log2 expression level varies from 4.5 to 5.3, or around 1.7 fold change, which is consistent with the fold change observed in *Gdnf*^wt/cHyper^;Pcp2-Cre animals (see Fig. 2B). This consistency supports the physiological relevance of our findings to normal human motor learning abilities.

The main inhibitory drive to Purkinje cells on the other hand is provided by basket cells and stellate cell interneurons of the cerebellar molecular layer, which use the vesicular γ-aminobutyric acid transporter (VGAT, Brown et al., 2019). Analysis of the inhibitory (VGAT) synapses onto Purkinje cells revealed no difference in the number of VGAT vesicles in RET-expressing basket cells interneurons (Figure S3C), nor in the number of VGAT-positive presynaptic vesicles measured on the Purkinje cell soma (Figure S3D), consistent with unchanged inhibitory input in our patch-clamp experiments at P60-P100 (Figure S3A, B).

Considering the increased excitatory drive on PCs in *Gdnf*^cHyper/cHyper^;Pcp2-Cre mice with no change in inhibitory inputs, we asked whether activity of the PCs, and, consequently, the cerebellum output signal, has changed. To address this, we registered spontaneous action potential firing of PCs in the cerebellum lobe VIII of the adult *Gdnf*^cHyper/cHyper^;Pcp2-Cre mice and their control littermates in cell-attached mode. We found that the spontaneous firing frequency of PCs in *Gdnf*^cHyper/cHyper^;Pcp2-Cre mice was significantly higher than in controls (Figure 4E, F).

### 3.6 Analysis of GDNF expression in normal human cerebellum

Gdnf mRNA levels correspond to GDNF protein expression levels in the brain and in the peripheral tissues and according to the previous reports regulate brain dopamine system function (Kopra et al, 2017; Kumar *et al*, 2015; Mätlik *et al*, 2022; Kopra *et al*, 2015; Olfat *et al*, 2023), reproductive organs development (Li *et al*, 2019), as well as the size, composition, and function of the enteric nervous system (Virtanen et al, 2024).Our results demonstrating the postnatal effect of about 2-fold increase in PCs’ GDNF mRNA levels on PC function and motor learning prompted us to explore the normal variation range in GDNF expression in the human cerebellum. For this, we extracted GDNF mRNA expression levels (Log_2_-transformed) from a publicly available dataset (GSE46706), from the study published by Ramasamy et al. (Ramasamy et al., 2014). The dataset contains expression data of the cerebellar cortex from 134 Caucasian healthy human individuals with a mean age at death of 59 years old (range 16–102). After filtering out extreme values of highest and lowest expression levels having low RNA Integrity Number (RIN<5) resulting in 120 samples, we found that the normal interindividual variation in GDNF expression levels in the human cerebellum is between 4.49 and 5.28 (logarithmic scale), which corresponds a fold change of 1.73 (Figure 6) and falls within the 2-fold variation range found in Pcp2-Cre GDNF Hypermorph mice relative to the control littermates (Figure 2B).

**Figure 6.**
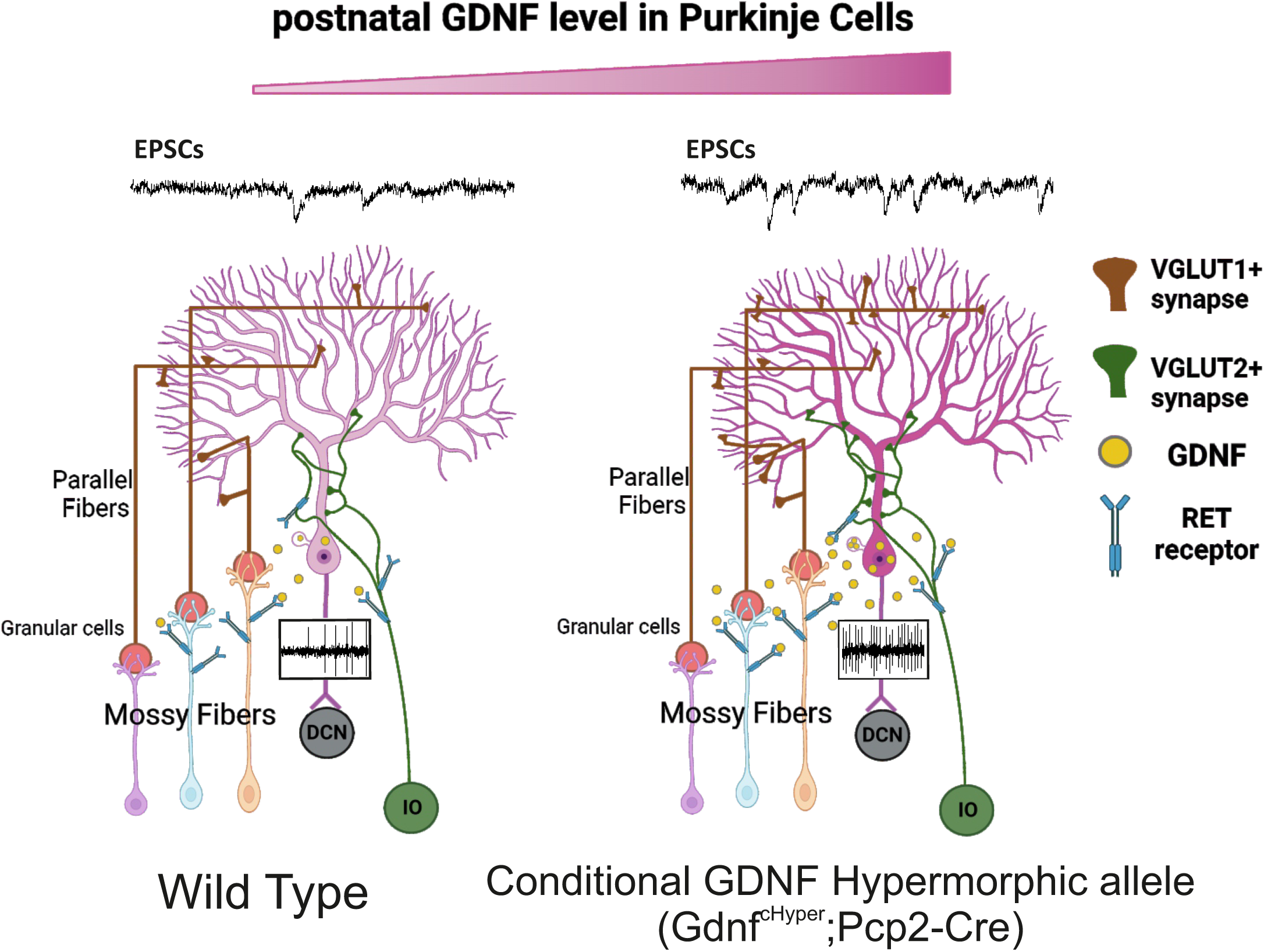
A model showing the effect of increased endogenous GDNF level in Purkinje cells. Elevated GDNF/RET signalling in Gdnf^cHyper^;Pcp2-Cre mice between GDNF-expressing Purkinje cells and RET-expressing glutamatergic axons of mossy fibers and Inferior Olives leads to an increased frequency of EPSCs in Purkinje cells (PCs, pink in the centre). This increase is likely due to both indirect upregulation of VGLUT1 synapses from Granular cells caused by RET-dependent changes in Mossy fibres and direct upregulation of VGLUT2 synapses from RET-expressing axons of Inferior Olives. The enhancement of excitatory drive onto PCs, without changes in inhibition (not shown at the scheme), increases the firing rate of PCs and overall output of the cerebellar cortex. These changes likely contribute to improved motor learning observed in Gdnf^cHyper^;Pcp2-Cre mice. Figure created with BioRender.com.

## 4 DISCUSSION

Understanding how specific brain functions are regulated by the expression levels of neurotrophic factors, in specific neuron types, has remained challenging. However, the ability to ask and answer such a question has broad relevance. For example, human GWAS-s show that about 93% of disease relevant as well as normal interindividual variation related differences lie within the non-coding genome (Edwards et al., 2013; Keaton et al., 2024; Tak and Farnham, 2015). Thus, interindividual variation including disease predisposition stems mainly from interindividual differences in gene expression regulation, not in their sequences. For some genes it is already well established that a small, 1,5-2-fold change in the final protein expression level can drive neurological disease (Liao *et al*, 2012; Elia *et al*, 2012; Glessner *et al*, 2009; Gennarino *et al*, 2015; Ross *et al*, 2008; Rovelet-Lecrux *et al*, 2006; Shao *et al*, 2021).

Conditional knock-out animals are useful experimental tools to elucidate gene function. However, often a 50% reduction in expression levels in heterozygous animals does not lead to a phenotype, whereas full gene deletion may be masked by molecular or cellular compensation, especially in the brain where alternative circuitries are known to be able to compensate for the hampered function (Grefkes and Fink, 2020; Kliemann et al., 2019; Moosa et al., 2013). Thus, hypermorph or overexpression alleles are often informative. GDNF itself serves as an example where results obtained from knock-out or conditional knock-out studies are most often not opposite or predictive to phenotypes observed in GDNF Hypermorph or conditional Hypermorph animals. For example, deletion of GDNF in the brain increases dopamine transporter activity (Kopra et al., 2017) but so does about 2-fold increase in endogenous GDNF levels (Kumar et al., 2015). The knock-out of GDNF encoding gene leads to a complete absence of the kidneys or the formation of miniscule kidneys (Moore et al., 1996). However, an increase in endogenous GDNF does not result in larger kidneys. Instead, it results in the formation of miniscule and malformed kidneys (Kumar et al., 2015) due to over-signalling at the ureteric bud tip cells (Li et al., 2019).

In this study, we used our novel conditional Gdnf Hypermorph mouse model (Mätlik et al., 2022) which allows Cre-driven upregulation of endogenous Gdnf at the post-transcriptional level, warranting increased expression only in natively transcribing cells. This model allowed us to investigate where, when, and how endogenous GDNF overexpression results in improved motor learning, previously reported in constitutive GDNF Hypermorph animals (Mätlik et al., 2018; Turconi et al., 2020). We first analysed the impact on the GDNF-expressing and GDNF responsive central motor control unit of the mammalian brain, the striatum, by injecting Cre-expressing virus specifically in this region of conditional GDNF Hypermorph animals. This manipulation increases striatal GDNF concentration by 2-3-fold in heterozygous animals (Figure 1E), and results in enhanced dopamine turnover (Mätlik et al., 2022; Olfat et al., 2023). Despite of this we did not observe changes in motor learning (Figure 1F). This suggested that GDNF levels are either important earlier in striatal development, influence other motor circuitries or act in the periphery such as the muscle tissue. Indeed, improved motor learning in *Gdnf*^wt/cHyper^;Nestin-Cre animals (Figure 1 C, D), with GDNF overexpression in the CNS starting from E12 argued for the CNS-mediated mechanism. Our Pcp2-Cre model showed that about 2 fold postnatal increase in GDNF expression specifically in Purkinje cells is sufficient to enhance motor learning seen in constitutive and Nestin-Cre-driven GDNF conditional Hypermorphs.

At the cellular level, we found that PCs in adult *Gdnf*^cHyper/cHyper^;Pcp2-Cre animals display higher frequency of spontaneous action potential firing. The opposite, reduction in frequency is reported in mouse models of spinocerebellar ataxia (SCA1-6, Cook et al., 2021), including SCA2 mouse model, where it was continuously decreasing with age starting from week 4 and accompanied by motor performance decline in the rotarod test (Hansen et al., 2013). Conversely, artificial increase of PCs’ action potential frequency with optogenetic stimulation has been shown to improve motor learning in several studies (Bonnan et al., 2021; Lindeman et al., 2020; Nguyen-Vu et al., 2013).

Our measurement of spontaneous post-synaptic currents and immunohistochemical analysis suggest that this increase in action potential firing of PCs can be caused by increased glutamatergic innervation from the climbing (VGLUT2+) and parallel (VGLUT1+) fibers. An alternative possibility is that elevated GDNF expression in PCs changes the intrinsic electrical properties of PCs, leading to an increased rate of action potentials. However, the well-established paracrine role of GDNF and the absence of RET on PCs (Figure 2F, S1, S4B, Sergaki and Ibáñez, 2017) make circuit-level changes a more reasonable explanation. (Airaksinen et al., 1999; Drinkut et al., 2016; Ibáñez and Andressoo, 2017; Taraviras et al., 1999). RET is expressed in inferior olive neurons, which give origin to VGLUT2-positive climbing fibers (Trupp et al., 1997), and GDNF is likely to upregulate their function through direct signalling and chemoattraction (Kumar et al., 2015; Love et al., 2005; Montaño-Rodriguez et al., 2024; Young et al., 2001). Another glutamatergic source onto PCs is coming from the granular cells (VGLUT1+), which in turn receive extensive glutamatergic input through mossy fibers from several nuclei in pons that also express RET (Trupp et al., 1997). Our experiments in *Ret*^GFP^ knock-in mouse line confirmed this result using IHC detection of GFP in the pontine nuclei (Figure S4A). We also found that some RET+ mossy fibers come in close proximity to PCs and may innervate at least some PCs (Figure S4B). The schematic summary of our findings is presented in Figure 6.

The transient, but very pronounce, communication between mossy fibers and PCs was shown during mouse postnatal development (Kalinovsky et al., 2011). The peak of this intercellular communication occurs at P7 and gradually declines by P21, followed by the formation of exclusive mossy fibers-to granule cells pathway. Considering that Pcp2-Cre-driven increase in endogenous GDNF expression begins in the first postnatal week, it is logical to suggest that during the first postnatal week, increased GDNF-RET signalling can affect the development of glutamatergic circuit within the cerebellar cortex.

Interestingly, we did not find changes in the number of the inhibitory RET-expressing basket cells, in their synaptic inputs or in their function on PCs in adult GDNF hypermorphs (Figure 2F, S1A-B, S3). Since our recordings in early postnatal constitutive GDNF Hypermorph mice showed a reduction in IPSCs in the PCs, one explanation is compensatory normalization by the time of the formation of the adult cerebellar circuit. However, longitudinal analysis on how the postnatal PCs-specific upregulation of GDNF expression affects the whole cerebellum circuit maturation in the development should be addressed in a separate study.

Our *Gdnf*^cHyper^;Pcp2-Cre mouse model demonstrated moderate fold change of the Gdnf mRNA expression in the cerebellar tissue exceeding the normal expression in control animals only up to about 2 times depending on allele dosage (Figure 2B). This level difference between control and PC-specific GDNF conditional Hypermorph animals is similar to that observed in healthy human adult cerebellum (Figure 5). Indeed, different interindividual abilities in motor learning are confirmed by a number of studies (reviewed in Anderson et al, 2021; Furuya, 2018). Based on our current results and results in mouse models where deletion or reduction in GDNF in the CNS (Kopra *et al*, 2015) or PCs (this study) does not affect motor performance it is feasible to speculate that while low or medium GDNF levels elicit no effect, more than average GDNF levels in the PCs may facilitate motor learning also in humans.

In conclusion, we show that endogenous GDNF levels in postnatal Purkinje cells is a non-essential but important regulator of motor learning. This effect is likely mediated through GDNF-RET signalling between Purkinje cells and glutamatergic cerebellum afferents.

## 5 AUTHOR CONTRIBUTIONS

E.N. and G.T. performed experiments, analysed and interpreted the data, prepared figures, and wrote the manuscript. G.T. and K.M. planned and designed experiments. K.M., M.S., S.O., E.N. performed experiments and analysed the data. E.N., M.S., T.T. interpreted the electrophysiology data and revised the manuscript, V.I. bred animals and revised the manuscript. J.O.A. planned and designed experiments, interpreted the data, provided funding and contributed to writing the manuscript. All authors read and approved the manuscript.

## 7 ACKNOWLEDGMENTS

G.T. was supported by the Finnish Parkinson’s foundation and the Doctoral School in Health Sciences. K.M. was supported by the Doctoral Programme Brain & Mind. E.N. was supported by Yrjö Jahnsson Foundation. J.O.A. was supported by the Academy of Finland (grants no. 297727 and 350678), Sigrid Juselius Foundation, ERA-NET NEURON grant nr 352077, Center of Innovative Medicine (CIMED), Hjärnfonden, Swedish Research Council (grants no. 2019-01578 and 2022-01093), Helsinki Institute of Life Science, and by European Research Council (ERC, grant no. 724922). The authors thank Prof. Chris de Zeeuw for the critical comments on the manuscript and Daniel R. Garton for language editing. Authors thank Biomedicum Imaging Unit and Mouse Behavioural Phenotyping Facility of the University of Helsinki.

---

**Supplementary Figure 1.**
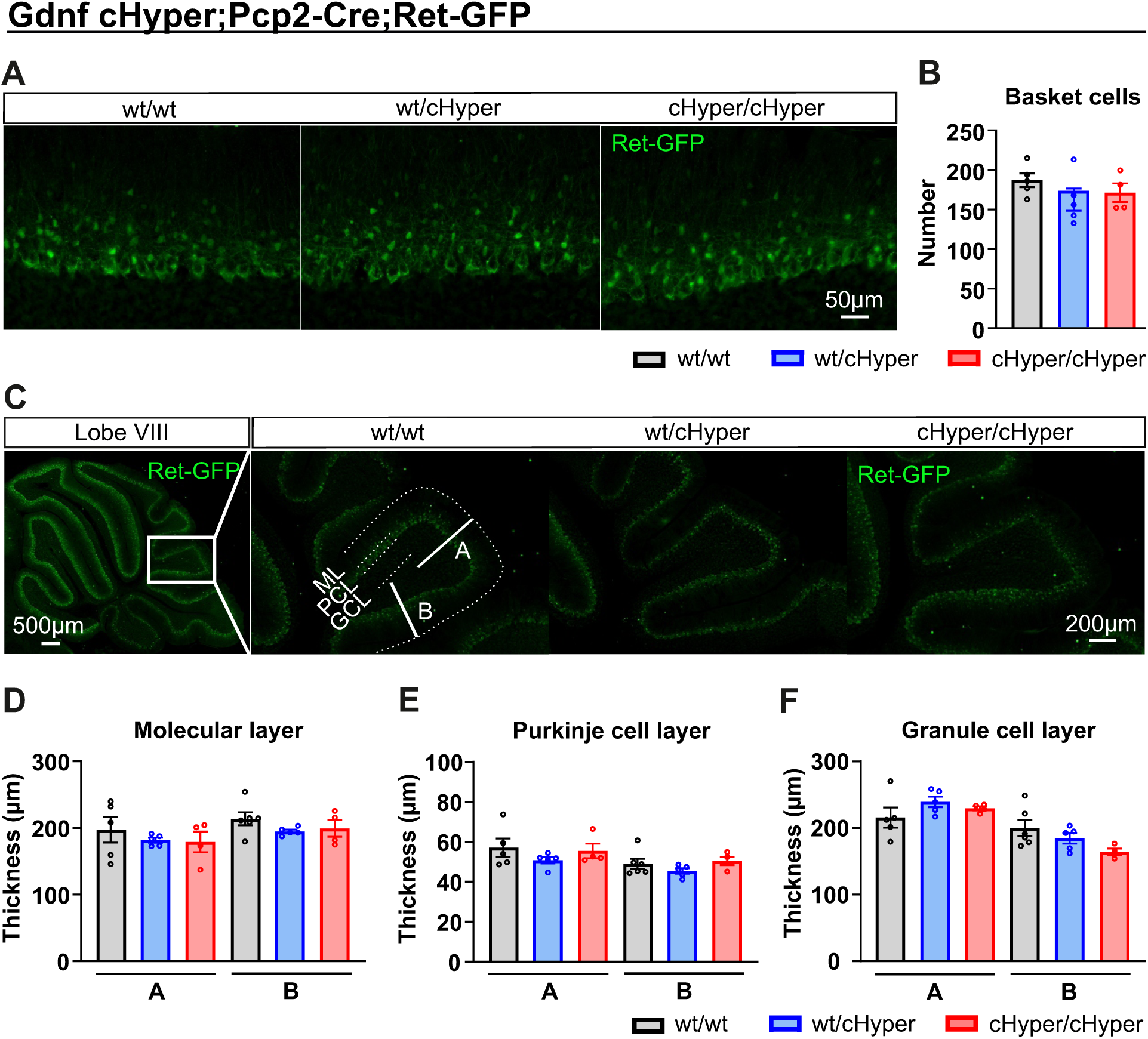
Unaltered number of RET-expressing cells and preserved cerebellar morphology upon endogenous GDNF increase in Purkinje cells. **(A)** Representative sagittal sections of cerebellar lobe VIII from adult *Ret*^GFP^ mice showing RET signal (green). **(B)** Number of RET-expressing basket cells in cerebellar lobe VIII. One-way ANOVA followed by Tukey’s multiple comparisons test. **(C)** Representative sagittal sections of cerebellar lobe VIII from adult *Ret*^GFP^ mice. The thickness of the molecular layer (ML), Purkinje cell layer (PCL) and granule cell layer (GCL) was measured at two different locations, named A and B. Scale bars are indicated in the figures. **(D-F)** Thickness of the cerebellar cortical layers measured at A and B locations from the Figure C in *Gdnf*^wt/wt^;Pcp2-Cre, *Gdnf*^wt/cHyper^;Pcp2-Cre, and *Gdnf*^cHyper/cHyper^;Pcp2-Cre mice. One-way ANOVA followed by Tukey’s multiple comparisons test. Data are presented as mean ± SEM. **(B, D-F)** *Gdnf*^wt/wt^;Pcp2-Cre (n=5); *Gdnf*^wt/cHyper^;Pcp2-Cre (n=5); *Gdnf*^cHyper/cHyper^;Pcp2-Cre (n=4).

**Supplementary Figure 2.**
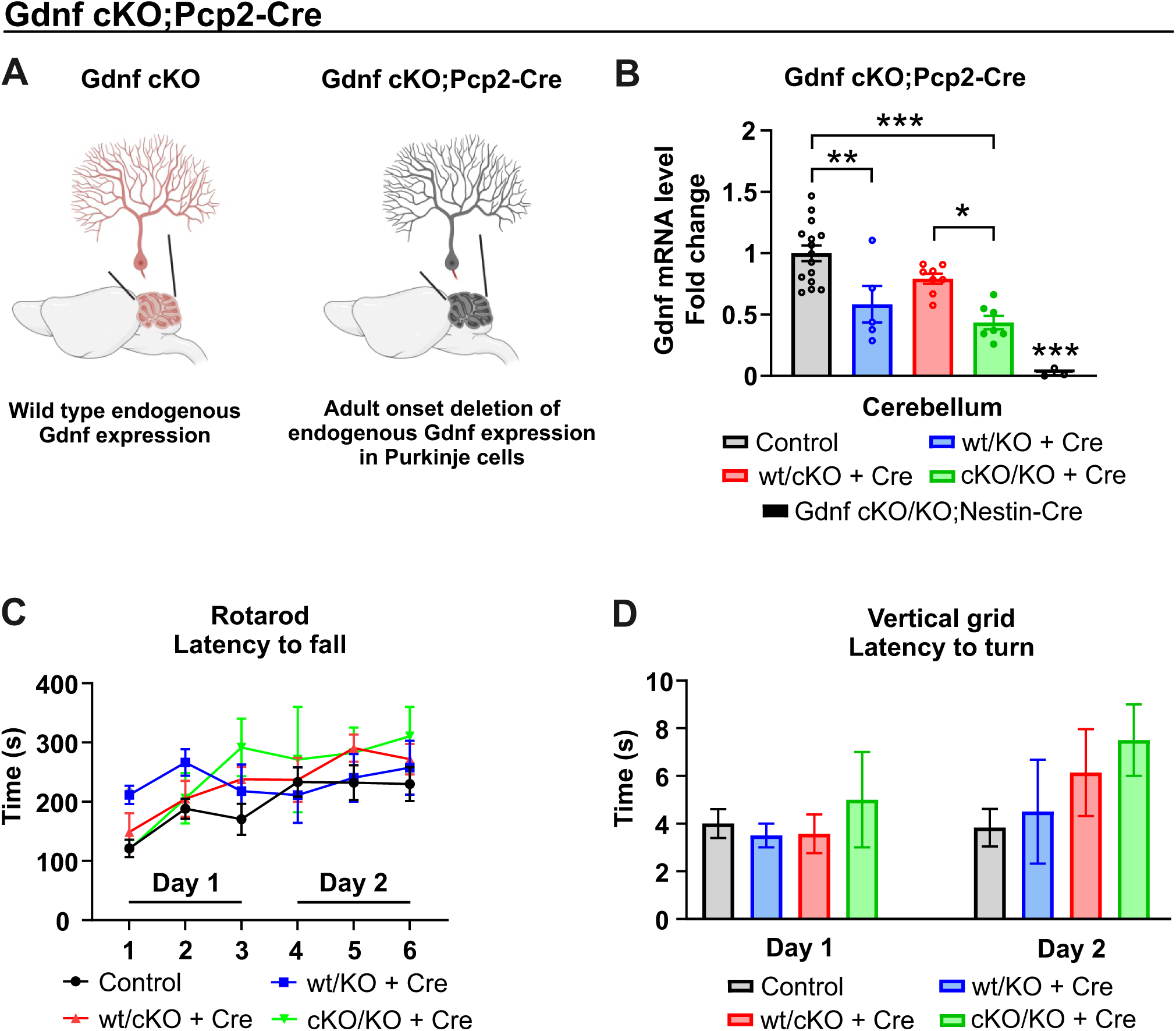
GDNF deletion in Purkinje cells does not alter motor function. **(A)** Endogenous GDNF deletion in cerebellar Purkinje cells was achieved by crossing the *Gdnf*^cKO^ allele with the Pcp2-Cre mouse line (*Gdnf*^cKO^;Pcp2-Cre). **(B)** Gdnf mRNA level in the adult cerebellum measured with qPCR. One-way ANOVA followed by Tukey’s multiple comparisons test. *p < 0.05, **p < 0.01, ***p < 0.001. **(C)** Latency to fall in the rotarod test and **(D)** latency to turn from the grid in the vertical grid test on experimental Day 1 and Day 2. Two-way ANOVA followed by Bonferroni’s multiple comparison test and unpaired t-test. Data are presented as mean ± SEM. **(B-D)** Control (n=12-15); *Gdnf*^wt/KO^;Pcp2-Cre (n=4-6); *Gdnf*^wt/cKO^;Pcp2-Cre (n=8); *Gdnf*^cKO/KO^;Pcp2-Cre (n=3-7); *Gdnf*^cKO/KO^;Nestin-Cre (n=3). Figure created with BioRender.com.

**Supplementary Figure 3.**
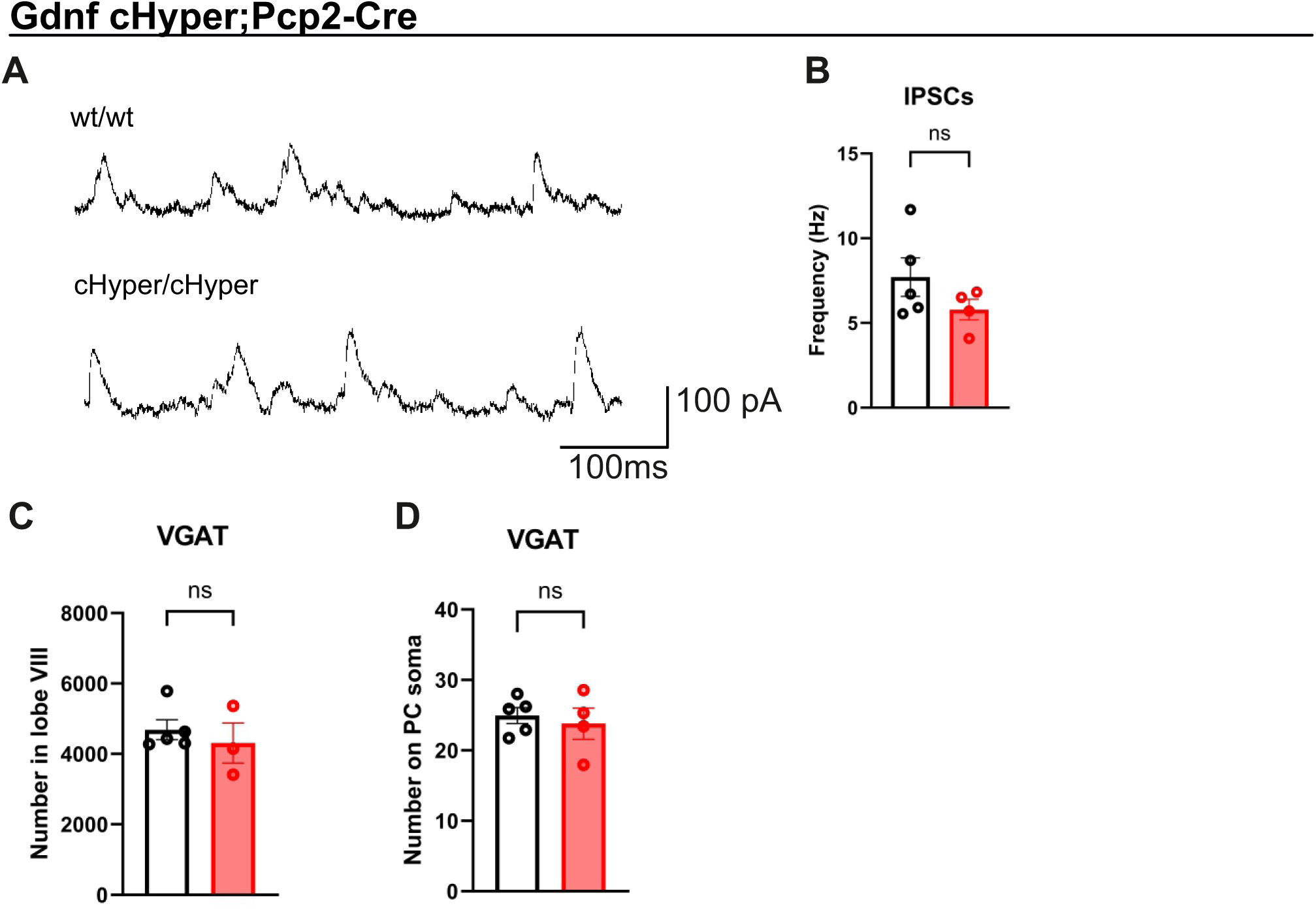
No changes in inhibitory synaptic inputs in Purkinje cells in *Gdnf*^cHyper^;Pcp2-Cre mice. **A)** Sample traces of spontaneous inhibitory post synaptic currents (IPSCs) in *Gdnf*^cHyper/cHyper^;Pcp2-Cre mice and their littermate controls. IPSCs are seen as outward currents (upward deflections) at the −40 mV holding potential. **(B)** Spontaneous IPSC frequency in Purkinje cells measured in cerebellar lobe VIII of *Gdnf*^wt/wt^;Pcp2-Cre adult mice and littermate controls. Unpaired t-test. **(C)** Number of vesicular GABA transporter (VGAT) synapses in 200 µm^2^ area within the PC layer. Unpaired t-test. Data are presented as mean ± SEM. **(B)** *Gdnf*^wt/wt^;Pcp2-Cre (n=5); *Gdnf*^cHyper/cHyper^;Pcp2-Cre (n=4). **(C)** *Gdnf*^wt/wt^;Pcp2-Cre (n=5); *Gdnf*^cHyper/cHyper^;Pcp2-Cre (n=3).

**Supplementary Figure 4.**
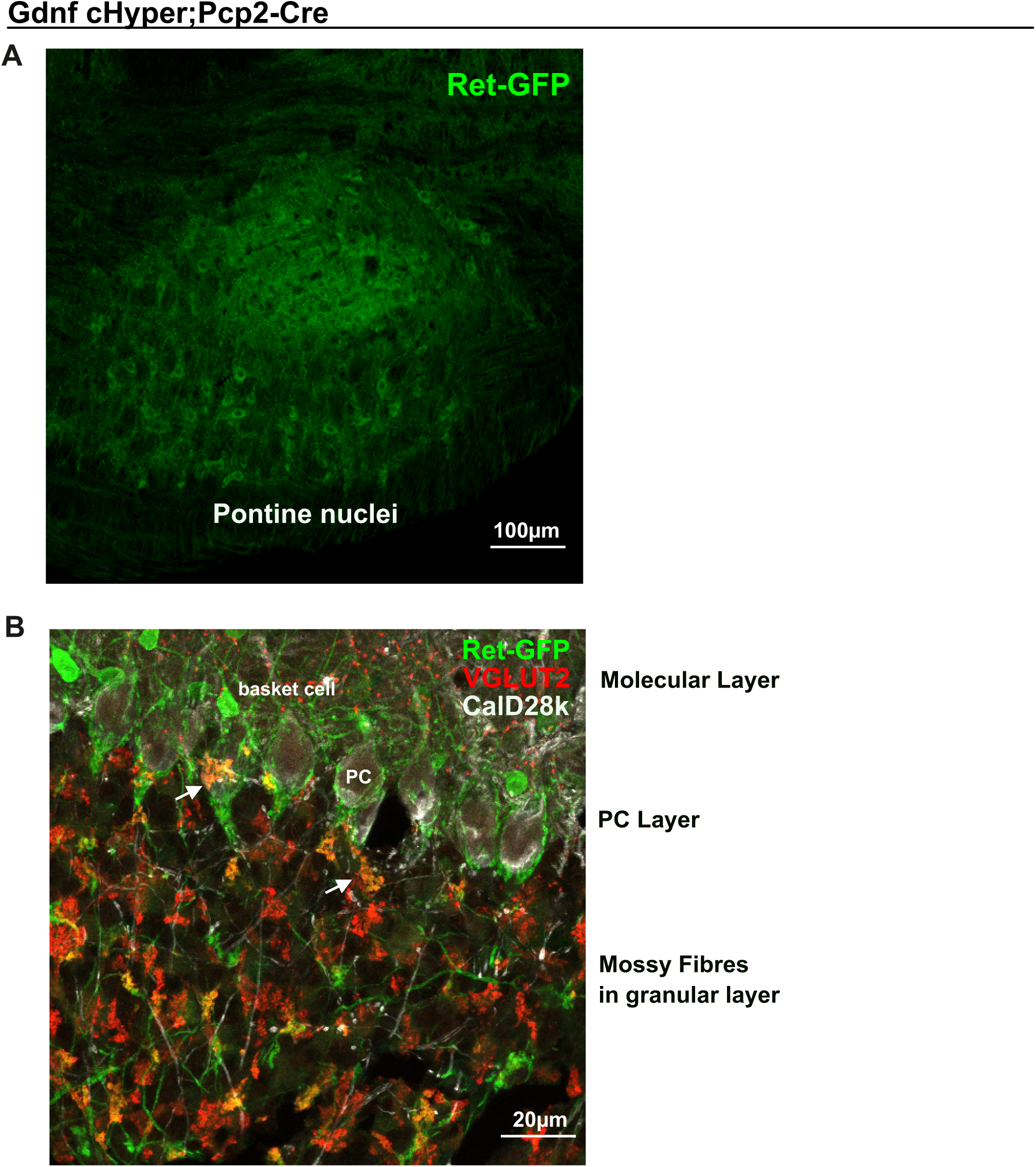
RET expression in knock-in *Ret*^GFP^ mouse brain. **(A)** Representative epifluorescent image of RET+ cells (green) in the pontine nucleus (sagittal plane) following anti-GFP staining. **(B)** Confocal image of the cerebellum lobe VIII region (sagittal plane), showing expression of RET (green), VGLUT2 (red) and CalD28k (white) close to the Purkinje Cells layer. White arrows indicate expression of RET in glutamatergic VGLUT2+ mossy fibres axons in the close proximity to PCs, suggesting direct GDNF signalling. Scale bars are indicated in the figures.

**Supplementary Table 1.** List of antibodies used in this study.

| **Primary antibody** | **Dilution** | **Company** | **Catalogue Number**<br>**#** |
| --- | --- | --- | --- |
| Rabbit anti-VGLUT2 | 1:500 | Synaptic Systems | 135 402 |
| Rabbit anti-VGAT | 1:1000 | Synaptic Systems | 131 002 |
| Mouse anti-CalD28K | 1:1000 | Swant | 300 |
| Goat anti-GFP | 1:1000 | Abcam | ab5450 |
| Rabbit anti-Parvalbumin | 1:1000 | Swant | pv27 |
| Goat anti-RFP | 1:500 | Rockland | 200-101-379 |
| **Secondary antibody** |  |  |  |
| anti-goat AlexaFluor 488 | 1:500 | Abcam | ab150129 |
| anti-goat AlexaFluor 594 | 1:500 | Abcam | ab150132 |
| anti-mouse AlexaFluor 488 | 1:500 | Abcam | ab150109 |
| anti-mouse AlexaFluor 594 | 1:500 | Abcam | ab150112 |
| anti-rabbit AlexaFluor 488 | 1:500 | Abcam | ab150073 |
| anti-rabbit AlexaFluor 594 | 1:500 | Abcam | ab150064 |

